# Levodopa’s effects on expression of reinforcement learning

**DOI:** 10.1101/434704

**Authors:** Grogan J.P., Isotalus H.K., Howat A., Irigoras Izagirre N., Knight L.E., Coulthard E.J.

## Abstract

Dopamine has been implicated in learning from rewards and punishment, and in the expression of this learning. However, many studies do not fully separate retrieval and decision mechanisms from learning and consolidation. Here, we investigated the effects of levodopa (dopamine precursor) on choice performance (isolated from learning or consolidation). We gave 31 healthy older adults 150mg of levodopa or placebo (double-blinded, randomised) 1 hour before testing them on stimuli they had learned the value of the previous day. We found that levodopa did not affect the overall accuracy of choices, nor the relative expression of positively or negatively reinforced values. This contradicts several studies and suggests that dopamine may not play a role in the choice performance for values learned through reinforcement learning in humans.

## INTRODUCTION

Dopamine has been heavily implicated in reinforcement learning^1–3^, and recently evidence has shown that dopamine also affects later choices based on these learned values^4–6^. However, unpicking the relative contribution of dopaminergic neurons during encoding, consolidation and retrieval stages of memory is often confounded by relatively long duration of action of medications.

### Exogenous dopamine administration biases consolidation or retrieval in Parkinson’s disease

An early study showed that if Parkinson’s disease (PD) patients were given their dopaminergic medication before completing a reinforcement learning task they learned better from positive than negative feedback^1^. The opposite pattern was shown if they were withdrawn from their dopaminergic medication prior to learning. However, the differences were not apparent during the learning trials themselves. Instead, after learning, all the combinations of stimuli were presented without feedback to see whether participants had learned the relative value of the symbols via positive or negative reinforcement. It was only on this latter choice test that the differences between medication states were seen, which raised the possibility that dopamine does not actually affect the learning process, but a separate process invoked when choosing stimuli based on their learned values. This could be a retrieval process for the learned values, or a decision process on the retrieved values.

When learning and choice trials were separated by a delay, which allowed PD patients to learn off medication and be tested on or off medication, medication state during learning had no effect on expression of positive or negative reinforcement, but dopaminergic state during the choices did^4^. This was accompanied by fMRI signals in the ventro-medial prefrontal cortex and nucleus accumbens tracking the value of stimuli only when PD patients were on medication. This suggested that dopamine improved the retrieval and comparison of the learned values.

Similarly, when PD patients learned a set of stimulus-stimulus associations, and only had the rewards mapped onto these stimuli after they had finished learning, they still showed a bias towards the most rewarded stimuli if they were on their medications during the entire session^5^. This demonstrated that the reward bias could be induced even when reward learning did not take place. Thus, dopamine appeared to affect value-based decision making, with a bias towards rewarding outcomes.

However, other studies have failed to find effects of dopamine during choice performance, with dopamine during testing 24 hours after reinforcement learning not affecting the change in accuracy from the learning trials^7,8^. One of these studies^7^ also found that PD patients on their dopaminergic medications during learning had poorer learning than those off medication. However, this task was a deterministic feedback task, rather than a probabilistic feedback task as used in most other studies, which may have different learning mechanisms due to the lack of stochasticity.

### Effect of dopamine administration in healthy young adults

While patients with Parkinson’s are known to be dopamine-depleted without medication, healthy young adults are usually considered to have optimal levels of dopaminergic activity for brain processing. Given the dopamine overdose hypothesis^9^ posits an optimal level of dopaminergic function, where both increases or decreases to this level impair functioning, one would predict distinct effects of dopamine administration on healthy young people compared to older people with relative dopaminergic loss^10^ and people with Parkinson’s disease who have more profound dopaminergic loss. Using the deterministic stimulus-response task mentioned above, healthy young participants were worse at learning after 100mg levodopa^11^. Likewise, pramipexole, a D2 agonist, impaired learning on the same task^12^. This could be explained by the increased dopaminergic activity tipping people over the peak of the inverted U-shaped response posited by the dopamine overdose hypothesis^13^.

A dopamine D2/3 receptor antagonist given to young adults during a probabilistic reward/punishment task did not affect the earlier stages of learning, but impaired performance at the later stages of the learning task, though only for the rewarded stimuli^6^. Computational modelling demonstrated an effect of dopamine on the choice parameter for the reward stimuli, but not for the punishment stimuli, or the learning rates, suggesting that the effect was not driven by learning from the feedback. This points to a D2/3 contribution to consolidation or retrieval of rewarded information in healthy young adults.

### Effects of exogenous dopamine in older adults

When healthy older participants were given levodopa before a reward/punishment learning task, they showed better performance on the reward trials, but no difference on the punishment trials, when compared against a haloperidol (D2 inverse agonist) group^14^. Neuroimaging revealed that levodopa increased the striatal reward prediction errors for reward trials but did not affect aversive prediction errors from the punishment trials. If contrasted with Eisenegger et al.^6^, it suggests that dopamine contributes to the reward prediction errors during learning, and that D2 receptors are important for the selection of actions, but not the learning from them. However, these studies used tasks with only learning trials, and used analysis techniques to try to separate out the influence of the drug on learning and choice selection within that.

Here, we used a separate choice phase on a reinforcement learning task which had no feedback, and thus tested choice selection only, to assess how levodopa affects the expression/retrieval of positive and negative learning. In order to isolate the effects of dopamine administration on choice performance from learning or consolidation, we gave this test phase 24 hours after initial learning and gave participants either 150mg levodopa or a placebo 1 hour before.

## METHODS

### Participants

Thirty-five healthy older adults were recruited from Join Dementia Research and the ReMemBr Group Healthy Volunteer database. One participant was excluded due to glaucoma (contraindication), and three withdrew before completing both conditions. Thirty-one participants completed both conditions.

Participants were native English speakers over 65 years old with normal or corrected vision. They had no neurological or psychiatric disorders and did not have any of the contraindications for the study drugs Domperidone and Madopar (levodopa; see Supplementary Materials 1). They were not taking any monoaminergic medications, or any drugs listed in the Summary of Product Characteristics for Domperidone or Madopar. Demographic details are provided in Table 1.

**Table 1.** The means, standard deviations (SD) and ranges of the demographic and questionnaire data for the participants.

| Measure | Mean | SD | Range |
| --- | --- | --- | --- |
| **N (Male: Female)** | 31 (14:17) |  |  |
| **Age** | 71.23 | 7.41 | 65 - 92 |
| **Years of Education** | 14.42 | 3.45 | 10 - 24 |
| **MoCA** | 26.19 | 3.10 | 18 - 30 |
| **DASS** | 11.29 | 10.12 | 1 - 39 |
| **BIS** | 57.53 | 9.01 | 38 - 73 |
| **LARS** | -26.65 | 5.45 | -34 - -14 |

Participants were tested at Southmead Hospital, Bristol, UK. All participants gave written informed consent at the start of each testing session, in accordance with the Declaration of Helsinki. Ethical approval was granted by University of Bristol Faculty Research Ethics Committee. All procedures were in accordance with Good Clinical Practice and HRA and ethical regulations.

### Design

A double-blinded, within-subjects, randomised placebo-controlled design was used. The two drugs were 10mg suspension of Domperidone and 187.5mg Madopar (37.5mg benserazide + 150mg levodopa) dispersible, both mixed with diluted squash, and the placebos were diluted squash, with a Vitamin C tablet dissolved in one to mimic the residue left by the Madopar dispersible tablet. The levodopa dose was chosen to match previous studies which have found effects of dopamine on reinforcement learning tasks ^15,16^.

Domperidone is a peripheral dopamine D2 receptor antagonist, given 1 hour before levodopa to counter the nausea sometimes caused by it. The drugs and placebos were prepared by a lab member not otherwise involved in the study.

### Tasks

The reinforcement task was adapted from Pessiglione et al.^14^, and is referred to as the GainLoss task. It was run using Matlab r2015 and Psychtoolbox-3^17–19^ on Dell Latitude 3340 laptops. Links to download the code are provided in the Data Availability section in this manuscript.

In this task, volunteers were instructed to attempt to win as much money as possible. During learning, on each trial one of three pairs of symbols was shown on the computer screen (Fig. 1). The participants selected one of them using the keyboard, after which their selection was circled in red for 500ms. This was followed by one of four outcomes presented on the screen for 1000ms: GAIN 20 pence; LOSE 20 pence; LOOK at a 20 pence piece; or NOTHING. The outcome was determined probabilistically, with symbol A in the Gain pair resulting in ‘GAIN’ on 80% of trials, and ‘NOTHING’ on 20%, and vice versa for symbol B in the Gain pair. In the Look pair, symbol C resulted in a ‘LOOK’ outcome 80% of the time, and ‘NOTHING’ 20% of the time (vice versa for symbol D), and in the Loss pair symbol F had an 80% chance of resulting in a ‘LOSS’ and 20% chance of ‘NOTHING’ (vice versa for symbol E). The outcome was displayed for 1000ms, which was followed by a fixation cross for 500ms before the onset of the next trial.

**Figure 1.**
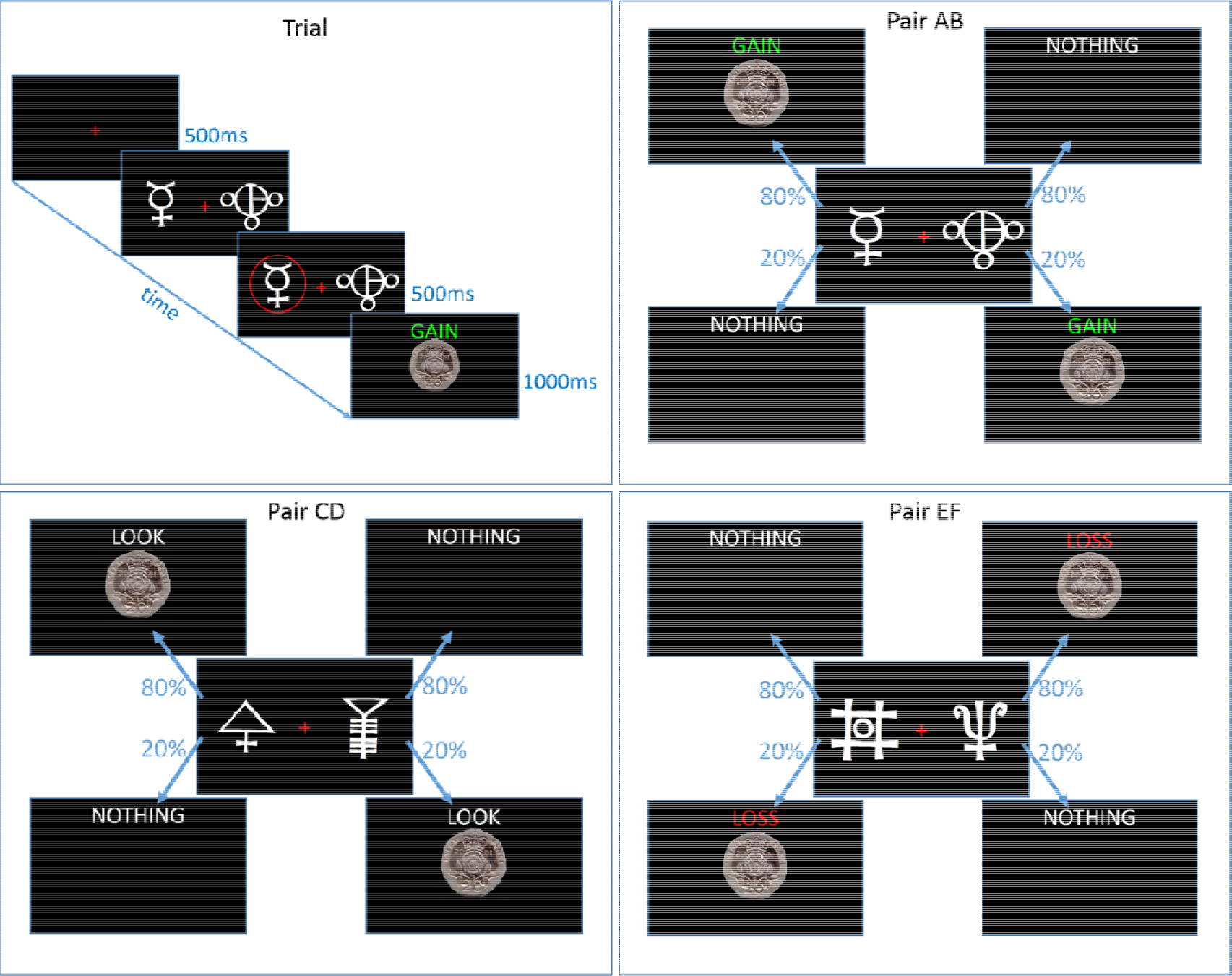
Diagram of the GainLoss experiment. Top left shows a sample Gain trial, and the other three panels show the outcome probabilities for the symbols in each pair.

The learning was preceded by a practice block of 30 trials (10 for each pair, using different symbols to the learning blocks), followed by two blocks of 90 learning trials (30 trials per pair). Choice performance was tested by showing all symbols in all combinations six times (e.g. AB, AC, AD…, 15 pairs in total, 6 repetitions of each pairs, 90 trials in total) without the outcomes shown. The stimuli were presented for the same duration as in the learning trials, except without the outcome screen. Choice performance was tested immediately after learning, after a 30-minute delay, and 24 hours later. Different sets of stimuli were used for each condition, the order of which was randomised across participants.

An episodic verbal learning task was also learned on day 1. Participants read aloud a list of 100 words and were tested 30 minutes and 24 hours later with the remember-know paradigm. Several questionnaires and paper tests were also given; digit span and the SMHSQ were given each day, and the MoCA, BIS, LARS, DASS and REI were given once each.

### Procedure

Participants completed four testing sessions, arranged into two pairs of days (see Fig. 2). On day 1, participants gave consent and were fully screened for all contraindications and interactions for the study drugs (Domperidone and Madopar), and Vitamin C, which was used in the placebo. They then learned the cognitive tasks and completed some of the questionnaires during the 30-minute delay before being tested on the tasks.

**Figure 2.**
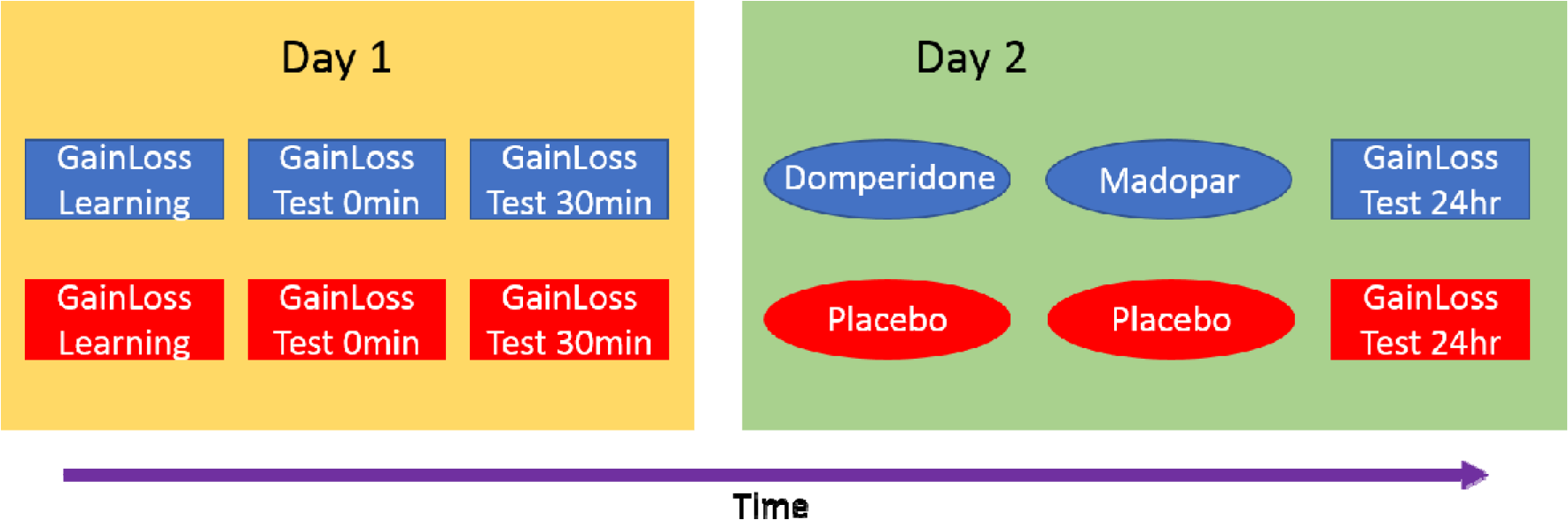
Timeline of experimental conditions. Each condition was identical except that in one pair of days participants received the drugs (blue) 1 hour before testing on Day 2, and on the other received the placebos (red) before testing. The order of drug and placebo condition was randomised across participants.

On day 2, participants again gave consent and continued eligibility was confirmed. Baseline blood pressure and heart rate was recorded before the Domperidone (or placebo; double-blinded) was administered. Thirty minutes later their blood pressure and heart rate were measured again, and the levodopa (or placebo) was given. Blood pressure and heart rate were also recorded 30 and 60 minutes later. One hour after the levodopa (or placebo) was administered, participants completed the GainLoss and remember-know tasks, digit span and SMHSQ. They then learned another list of words to test encoding effects of dopamine on long term memory, and memory was tested immediately, and over the phone 1, 3 and 5 days later.

Days 3 and 4 were identical to days 1 and 2, with the exception of the drug/placebo. On the last phone test after day 4, participants were asked which day they thought they received the drugs to assess blinding success.

### Data analysis

Selection of the symbol that was more likely to lead to the highest value of the two shown was considered the optimal response, regardless of the outcome actually given on that learning trial (e.g. if they select symbol A, the 80% Gain symbol, this is considered optimal even it results in ‘NOTHING’ on that particular trial). For the Look pair, symbol C (80% LOOK) was treated as optimal when it was against ‘NOTHING’ even though neither outcome had monetary value. The Look symbols were considered optimal against the Loss symbols, while the Gain symbols were considered optimal against the Look symbols.

For the choice tests, the number of times each symbol was chosen was divided by the number of times it was seen, to give percentage selections (see Fig. 3). Percentage avoidances were calculated likewise. Within-subject ANOVAs and t-tests were used on the 24-hour test measures to see how levodopa affected choice performance. Cohen’s *d* and partial *η*^*2*^ ( ) effect sizes are reported alongside t-tests and ANOVAs, respectively. We used SPSS v23 (IBM) for statistics. Q-Q plots were used to verify that data were approximately normal before parametric tests.

**Figure 3.**
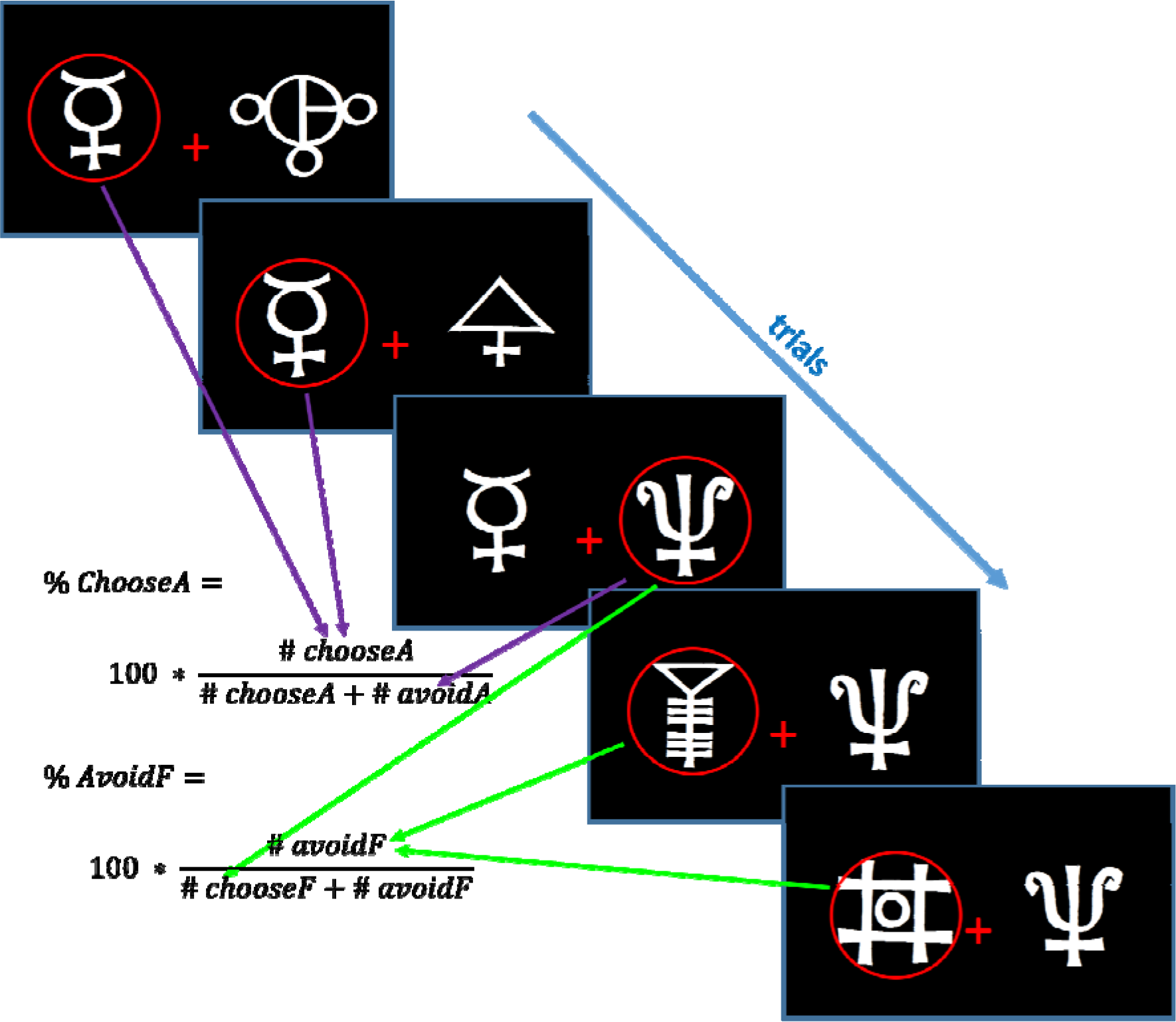
Diagram showing how Choose-A and Avoid-F were calculated. The same procedure was used for all symbols.

In addition to frequentists statistical analyses, we also performed Bayesian analyses in JASP^20^. Bayesian t-tests and repeated measures ANOVAs were used. Bayesian analysis compares the likelihood of the data given the null hypothesis (H_0_) to the likelihood given the experimental hypothesis (H_1_). The ratio of these two gives the Bayes Factor (BF_01_ = H_0_/H_1_) which quantifies how much more likely the data are given the null hypothesis rather than the experimental hypothesis. Please note that BF can also be reported in terms of the experimental hypothesis (i.e. BF_10_ = H_1_/H_0_), but we use the BF_01_ here due to the direction of results we found. BF of 1 suggest equal evidence for the two hypotheses, while the further the BF is from 1, the stronger the evidence for or against the null. We used the default prior of a Cauchy distribution with width 0.707 (meaning we assume there is a 50% probability of the effect size being between −0.707 and 0.707). Robustness checks with different prior widths are provided in the Supplementary Materials.

A previous study showed dose-dependent effects of levodopa on episodic memory consolidation when body weight was used to adjust the doses^21^. Body weight affects total absorption of levodopa, and the elimination half-life^22^. Therefore, we divided the levodopa dose (150mg) by body weight (kg) to give the weight-adjusted doses (mg/kg) and looked for linear or polynomial regressions between this and the difference in accuracy and choices between drug and placebo conditions.

## RESULTS

Participants were not able to guess correctly which day they received the drugs or placebo. Twenty-nine participants provided guesses, of which 17 were correct, and a binomial test showed this was not significantly different from chance (p = .720).

### Accuracy

During learning trials, participants performed best on the Gain pair (mean 57% accuracy, SD = 14.4), and worse on the Loss pair (mean = 53%, SD = 11.8; Look pair mean = 52% %, SD = 14.0). However, the mean accuracies were much higher for the choice tests at 0 minutes, 30 minutes and 24 hours (>65%; see Fig. 4). As the drug/placebo was only given before the 24-hour test, we used paired t-tests to look at the accuracy on this test, which revealed accuracy was not affected by levodopa (t (30) = 0.906, p = .372, *d* = .163; BF_01_ = 3.581).

**Figure 4.**
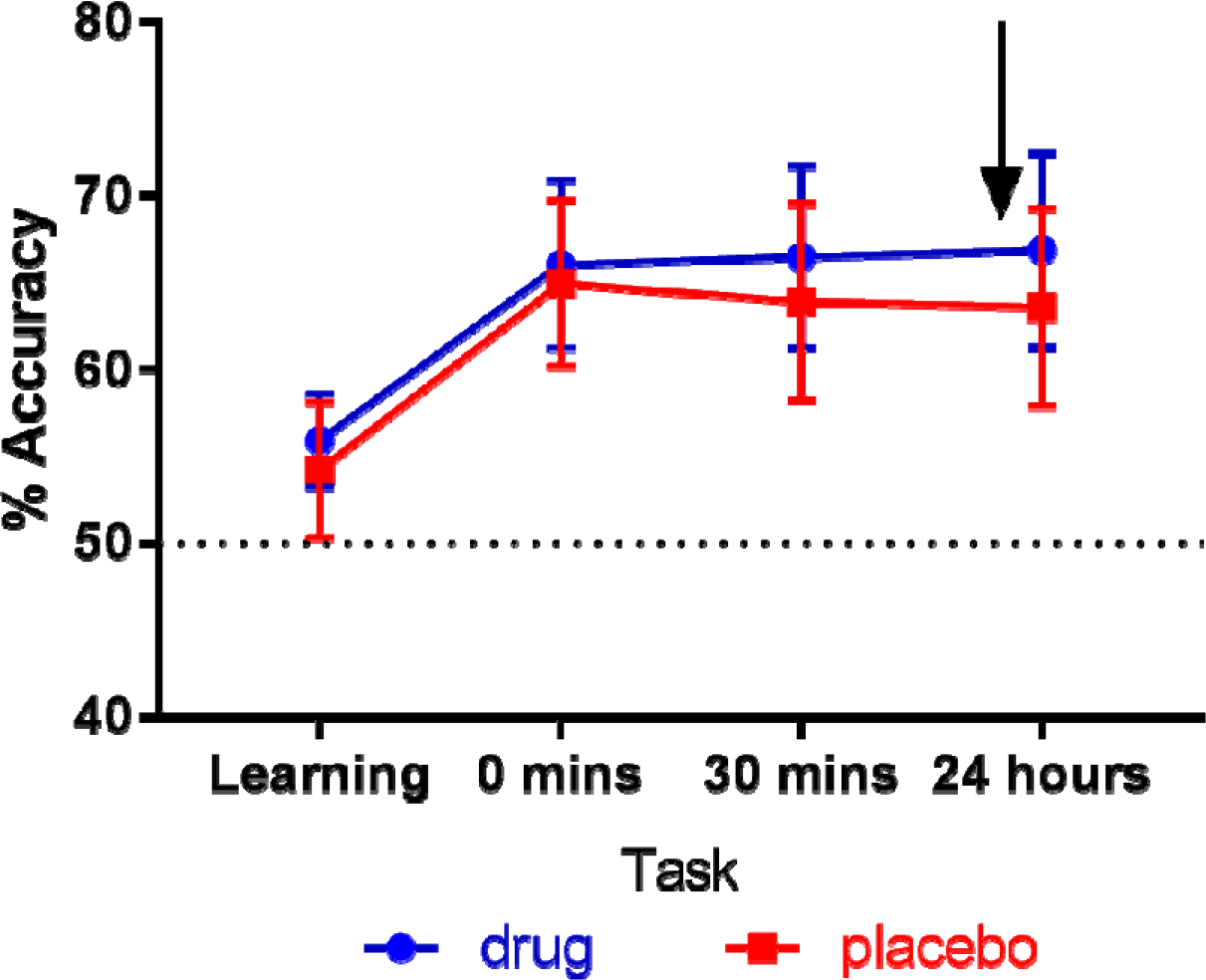
The mean % accuracy on learning and novel pairs tests, for both conditions (95% confidence intervals). The arrow shows when the drug/placebo was administered (time not to scale). There was no difference between accuracy after drug or placebo (p = .372, BF_01_ = 3.581).

### Positive and negative choices

We divided the number of times participants choose each symbol by the number of times it was presented to give the percentage of choices of each symbol (see Fig. 3). Figure 5 shows the mean percentages of the choice of each symbol for the drug and placebo conditions at 24 hours. Paired t-tests showed no significant differences in percentage of choices on drug or placebo for any of the symbols (p > .05; see Table 2 for statistics), suggesting that levodopa did not affect selection for any choice. Bayesian t-tests showed moderate evidence in favour of the null hypothesis (BF_01_ > 3) for all choices apart from symbol F where the evidence for the null hypothesis was anecdotal (BF_01_ = 1.301; see Table 2). This suggests that levodopa does not affect choice selection, except for the most punished symbol where the evidence is inconclusive.

**Table 2.** Statistics from frequentist and Bayesian t-tests on the accuracy and percentage of choices for each symbol at the 24-hour choice test. BF_01_ > 3 reflects moderate evidence in favour of the null hypothesis. Cohen’s d and 95% confidence intervals are presented for frequentist t-tests, and the posterior median and 95% credible intervals for the Bayesian t-tests. All error % from the Bayesian analyses were < 4×10^−4^.

| Measure | t | p | d | 95% Conf Int | $BF_{01}$ | Posterior | 95% Cred Int |
| --- | --- | --- | --- | --- | --- | --- | --- |
| **Accuracy** | 0.906 | .372 | .163 | -4.135, 10.730 | 3.581 | 0.148 | -0.186, 0.482 |
| **Choose-A** | -0.332 | .742 | -.060 | -16.919, 12.188 | 4.960 | -0.052 | -0.391, 0.281 |
| **Choose-B** | 0.718 | .478 | .129 | -8.725, 18.187 | 4.115 | 0.115 | -0.214, 0.455 |
| **Choose-C** | 0.878 | .387 | .158 | -6.981, 17.519 | 3.663 | 0.143 | -0.200, 0.494 |
| **Choose-D** | 0.108 | .915 | .019 | -13.463, 14.968 | 5.192 | 0.019 | -0.308, 0.355 |
| **Choose-E** | 0.454 | .653 | .082 | -11.277, 17.729 | 4.744 | 0.071 | -0.252, 0.415 |
| **Choose-F** | -1.771 | .087 | -.318 | -25.002, 1.177 | 1.301 | -0.288 | -0.642, 0.055 |

**Figure 5.**
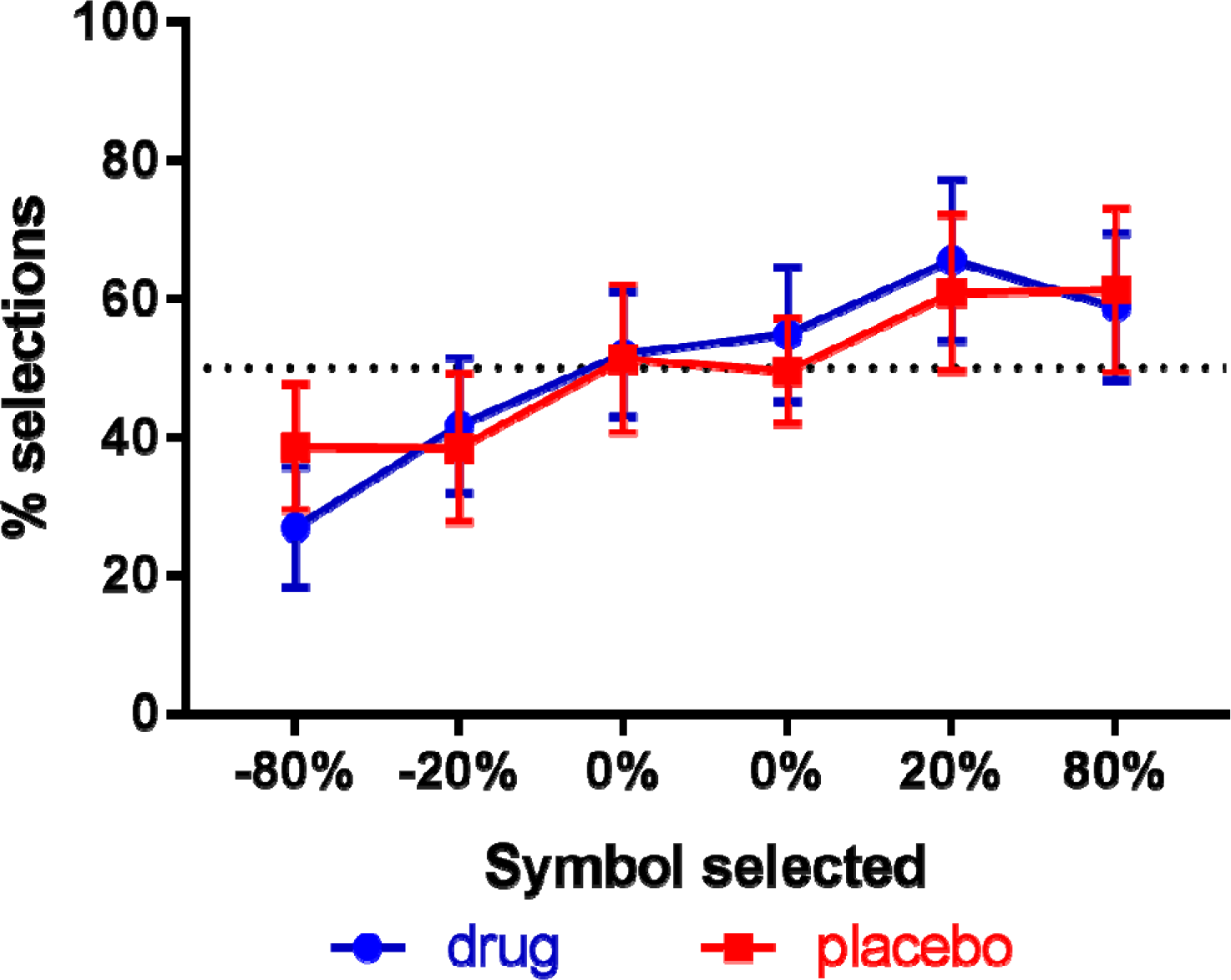
The mean percentage of choices of each symbol in the 24-hour choice test for both conditions (95% confidence intervals). The value of the symbol is the sum of the probability multiplied by the value of each outcome (i.e. 80% chance of loss (−1) and 20% chance of nothing (0) gives −80%). There were no significant differences between drug and placebo conditions for any symbol (p > .05, BF_01_ > 1).

We ran a repeated measures ANOVA to see whether levodopa affected the selection of the most rewarded and punished symbols differently (this is analogous to the ANOVAs run on choose-A and avoid-B in Frank et al.^1^). Looking just at the number of times the most rewarded symbol was chosen (choose-A) and the number of times the most punished symbol was avoided (avoid-F), there was no effect of medication (F (1, 30) = 0.719, p = .403, 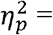 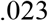) nor an interaction of medication and choice (F (1, 30) = 2.851, p = .102, 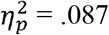). This again suggests that levodopa did not affect expression of positive or negative reinforcement (Fig. 4; avoid-F is the inverse of choose-F). A Bayesian repeated measures ANOVA found that this data was most likely under the null model (with no effects of medication, choice, or interactions; BF_M_ = 3.252) arguing against the inclusion of medication or choice in the model (BF_inclusion_ < 1).

Weight-adjusted dose did not have any significant linear or polynomial associations with the difference (between levodopa and placebo conditions) in 24-hour choice accuracy or on the difference on any of the choices (*r*^*2*^ < .017, p > .2). Nor did we find any associations between 24-hour accuracy or choice behaviour and MoCA, DASS, BIS, LARS, age, or years of education (p > .05; see Supplementary Materials). Four participants had low MoCAs (<24), so we reran the main analyses excluding these to see if the lack of results was due to cognitively impaired participants reducing effects. The pattern and significance of results was no different to the analyses presented above (see Supplementary Materials).

## DISCUSSION

Levodopa given 24 hours after learning a reward and punishment task did not affect choice performance. This suggests that levodopa, and hence dopamine, does not affect the expression of positive or negative reinforcement 24 hours after learning.

This contradicts several other studies which have found that dopamine can affect expression of reinforcement learning^4–6^. However, there are several differences between each of these studies and the current one. For example, Shiner et al.^4^ and Smittenaar et al.^5^ did not have punishments in their task, only rewards of varying probabilities. It may be that dopamine’s effects are only seen on positive reinforcement, which were missed in our task as we only had 2 stimuli that were positively reinforced (symbols A and B).

Eisenegger et al.^6^ used a task with positive and negative reinforcement like ours but did not have a separate ‘novel pairs’ choice test. Instead, they looked at the performance towards the end of the learning trials and used that to assess effects on the expression of learning. While their modelling analysis suggested the effects were not due to differences in learning rates, but rather the softmax decision parameter, this was still during the learning process and thus may be quite different to processes that occur much later and do not incur feedback. It should be noted that the softmax parameter in reinforcement learning models captures how frequently participants make a ‘greedy’ selection and choose the stimuli with the highest value, rather than making an explorative choice to a lower value stimulus. Thus, it also functions like a noise parameter, and will be higher when there is more variance that the learning rate parameters cannot explain. It is possible that the true effects were not due to more random choosing but rather some unknown process during learning that was simply captured by this noise parameter.

Alternatively, perhaps our participants did not learn the task well enough for us to be able to detect differences. The average accuracy at the end of the learning trials was close to chance, though it increased on the novel pairs tests to levels seen in other studies^1,4,5^ (i.e. 50-80%). The poor learning may have been compounded by the inclusion of several participants with low MoCAs, although excluding these participants did not change the pattern or significance of results.

The lack of effect of levodopa on anything could also suggest that the drug simply was not having an effect. However, we used a fairly large dose (150mg levodopa), which is as large as or larger than several other studies^11,14–16,23–25^. We waited 1 hour between dosing and testing, which coincides with the time to max concentration^26,27^. Some studies have reported dose-dependent effects, with people who had larger relative doses (due to smaller body weight, thus higher mg/kg dosage) showing greater effects^15,22^. We found no such associations, linear or quadratic. This suggests the lack of effect was not due to the specific dosage given.

Finally, if dopamine does not affect expression of reinforcement learning, then how can we explain previous results? Consolidation is a mechanism often overlooked in this type of memory, but in between learning the values and retrieving them, those values must be stored for a period and protected against interference from other learning. It may be that previous effects can be explained by consolidation, as the dopamine drugs were either present during learning (and thus consolidation after learning) or given just after learning before a 1-hour delay (which would have allowed consolidation to be affected). Previous studies have suggested dopamine may affect the persistence of reinforcement learning across time^8,28^, and this is a possible avenue for future research.

## ACKNOWLEDGEMENTS

The authors wish to thank NBT and BRACE for the use of their buildings, the funders (Wellcome [097081/Z/11/Z], MRC and BRACE [14/8108]), and all the participants who took part in the study.

## AUTHOR CONTRIBUTIONS

JPG, HKI and EJC designed the study. JPG, HKI, LEK, AH and NII performed data collection. JPG analysed the data. JPG, HKI and EJC wrote the manuscript. All authors reviewed the manuscript.

## DATA AVAILABILITY

Data are available at the University of Bristol data repository, data.bris, at https://doi.org/10.5523/bris.qpqzeqc3q53m2dwczp69q3pv0 ^29^. Our Matlab code for the analysis is available here https://doi.org/10.5281/zenodo.1438407 ^30^, and the code for the GainLoss task is available here https://doi.org/10.5281/zenodo.1443384 ^31^.

## ABBREVIATIONS

BIS: Barratt Impulsivity Scale
DASS: Depression, Anxiety and Stress Scale
LARS: Lille Apathy Rating Scale
MoCA: Montreal Cognitive Assessment
PD: Parkinson’s disease

